# Attractive internuclear force drives the collective behavior of nuclear arrays in *Drosophila* embryos

**DOI:** 10.1101/2020.10.08.330845

**Authors:** Xiaoxuan Wu, Kakit Kong, Wenlei Xiao, Feng Liu

**Affiliations:** Center for Quantitative Biology, Peking University, China; State Key Laboratory of Nuclear Physics and Technology, School of Physics, Peking University, China; School of Mechanical Engineering and Automation, Beihang University, China

**Author notes:** Correspondence (F.L.).

**Keywords:** collective behavior, *Drosophila* embryo, nuclear array, deep neural network, force field

## Abstract

The emerging collective behaviors during embryogenesis play an important role in precise and reproducible morphogenesis. An important question in the study of collective behavior is what rule underlies the emerging pattern. Here we use the *Drosophila* embryo as a test tube to study this question. We focus on the nuclear array without membrane separation on the embryo periphery from the nuclear cycle (NC) 11 to NC14. After live imaging with light sheet microscopy, we extract the nuclear trajectory, speed, and internuclear distance with an automatic nuclear tracing method. We find that the nuclear speed shows a period of standing waves along the anterior-posterior (AP) axis after each metaphase as the nuclei collectively migrate towards the embryo poles and partially move back. And the maximum nuclear speed dampens by 38% in the second half of the standing wave. Moreover, the nuclear density is 35% higher in the middle than the pole region of the embryo during the S phase of NC11-NC14. To find mechanical rules controlling the collective motion and packing patterns of the nuclear array, we use the deep neural network (DNN) to learn the force field from data. We find two potential strong nuclear-age-dependent force fields, i.e., the repulsive or attractive force field. Simulations with the particle-based model indicate that only if the net internuclear force is attractive and increases with distance, the pseudo-synchronous mitotic wave in a nuclear array with lower nuclear density in embryo poles can drive the collective motion with the damped standing wave of the nuclear speed, and the collective nuclear motion, in turn, maintains the non-uniform nuclear density.

## INTRODUCTION

The emerging collective behaviors during embryogenesis are key to understand the origin of the precise and reproducible morphogenesis (Streichan et al., 2018, Deneke et al., 2019, Lv et al., 2020). In general, complex collective behaviors (such as cytoskeleton filament arrangement, morphogenesis, and bird flocks) arise from interactions between many similar units (such as cytoskeleton macromolecules, cells, and birds) (Vicsek and Zafeiris, 2012). To understand the mechanism underlying the collective patterns, it is essential to uncover the interaction rules between individual units (Lukeman et al., 2010, Katz et al., 2011, Hinz and de Polavieja, 2017, Merkel and Manning, 2017, Ishihara and Sugimura, 2012). However, it has been challenging to reversely infer the interaction rule from the collective motion data.

The nuclear array in the early embryo of *Drosophila melanogaster* is an excellent model system to address this question. During the first 13 nuclear cycles (NCs), the whole embryo is a synplasm, in which nuclei share a common cytoplasm and the corresponding molecular environment (Foe and Alberts, 1983, Farrell and O’Farrell, 2014). From NC10 to NC14, the nuclei distribute near the embryo surface to form a two-dimensional (2D) nuclear array (Foe and Alberts, 1983, Farrell and O’Farrell, 2014). After each nuclear cycle, the density of the nuclear array doubles, and the internuclear distance decreases. The spatial and orientation order increase from early to later nuclear cycles (Kaiser et al., 2018), but the radial distribution functions overlap if they are rescaled with the nuclear density (Dutta et al., 2019). At the end of each nuclear cycle, a persuade-synchronized mitotic wave usually starts from the two embryo poles, moves towards the equator of the embryo along the anterior-posterior (AP) axis. A collective “yo-yo”-like nuclear movement follows spindle splitting, i.e., the nuclei move towards the two poles then back nearly to the original position (Lv et al., 2020).

However, the mechanism controlling these patterns remains to be elucidated. An essential reason is the underlying complicated molecular dynamics. During the S phase, a nucleus is surrounded by microtubules and has an Arp2/3 nucleated actin cap attached to the membrane (Zhang et al., 2018, di Pietro and Bellaïche, 2018, Schmidt and Grosshans, 2018, Foe et al., 2000). In the internuclear region, myosin II and linear actin filaments form an actomyosin complex adhering to the membrane (Zhang et al., 2018, di Pietro and Bellaïche, 2018, Schmidt and Grosshans, 2018, Foe et al., 2000). During the M phase, the actin cap region enlarges, and the internuclear membrane invaginates to form mitotic furrows surrounding the spindles (Zhang et al., 2018, di Pietro and Bellaïche, 2018, Schmidt and Grosshans, 2018, Foe et al., 2000). These two states alternate over the repeated nuclear cycles.

The net effective interaction force that underlying the nuclear array could be either attractive or repulsive. If microtubules and the attached motors (e.g., kinesin-5) are the predominant factors for the internuclear force, the force would be repulsive (Fig. S7A) (Manhart et al., 2018, Kaiser et al., 2018, Mann and Wadsworth, 2019). The repulsive force could also originate from the passive response of the elastic nuclear-embedding cortex (Doubrovinski et al., 2017). If actomyosin borders are the predominant factors for the internuclear force, the force would be attractive (Fig. S7B) (He et al., 2016, Royou et al., 2002, Izquierdo et al., 2018, Deneke et al., 2019, Streichan et al., 2018, Foe et al., 2000). Microtubules attached with the motor dynein may also provide attractive force (Manhart et al., 2018). Microtubules interact with actin caps via dynein-dynactin complexes (Kanesaki et al., 2011, Robinson et al., 1999, Cytrynbaum et al., 2005), and in the absence of actin caps, adjacent nuclei collide (Kanesaki et al., 2011).

Several particle-based models have been proposed with different formulas of the pair-wise force between nuclei (Kaiser et al., 2018, Dutta et al., 2019, Lv et al., 2020). And nearly all of these models assume that the predominant internuclear force is repulsive. Through parameter fitting, each model appears to be attributed to the specific feature of interest. However, a coherent force field has yet to be tested to account for all the observed characteristic features of the nuclear array.

Inspiring by the idea “from data to rules”, we use the deep neural network (DNN) to directly learn internuclear interaction rules from the experimental data. This new approach has several advantages. First, without any arbitrary assumption of the specific formula, the learned rules could be unbiased to show the integrated effect of all of the related molecules. Second, since DNN is a universal approximation to any Borel measurable function with the desired degree of accuracy (Hornik et al., 1989, Svozil et al., 1997), the potential complex rules can be accurately represented. Last, benefitting from the development of TensorFlow, the fitting process is really efficient. With the input of the nuclear age and nuclear density, the network outputs the internuclear force. The training error is calculated according to the resultant force and nuclear speed. Through learning, we find that the force field is strongly dependent on the nuclear age. Simulations with the particle-based model indicate that the attractive force field is consistent with experimental results.

## RESULTS

### Nuclear array shows a stereotypic packing pattern and collective motion pattern from NC11 to NC14

To quantify the packing and collective motion patterns of the nuclear array in *Drosophila* embryos, we image His-GFP expressing embryos from NC10 to early NC14 with light sheet microscopy, which has low phototoxicity and high temporal resolution (Fig. 1). With an automatic segmentation and tracking algorithm based on TGMM (Tracking with Gaussian Mixture Models) (Yue et al., 2020, Amat et al., 2014), we obtain the nuclear position and speed (Fig. 1 and movie S1). We discover that the nuclear array shows a specific packing pattern and collective motion pattern during the early developmental process, and these patterns are conservative across embryos (Fig. 2 and Figs. S1-S6).

**Fig. 1.**
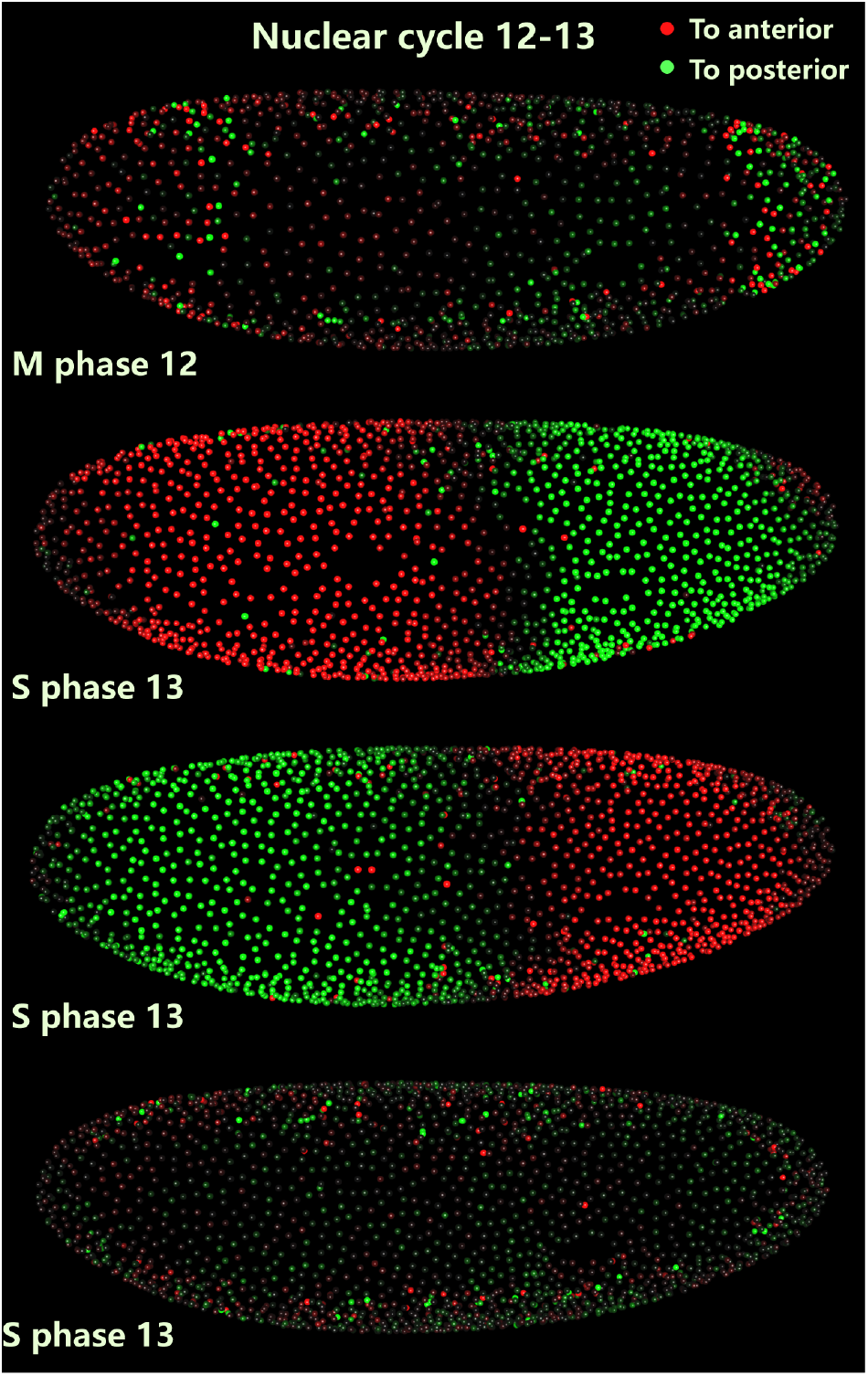
Representative images of the collective motion of the nuclear array of a *Drosophila* embryo (movie see movie S1). These images are the projections from the 3D images taken with light sheet microscopy. Each nucleus is segmented and marked with a different color to show the direction of the nuclear velocity along the AP axis (red and green represent the left (anterior) and right (posterior) direction, respectively). And the intensity indicates the magnitude of the nuclear velocity.

**Fig. 2.**
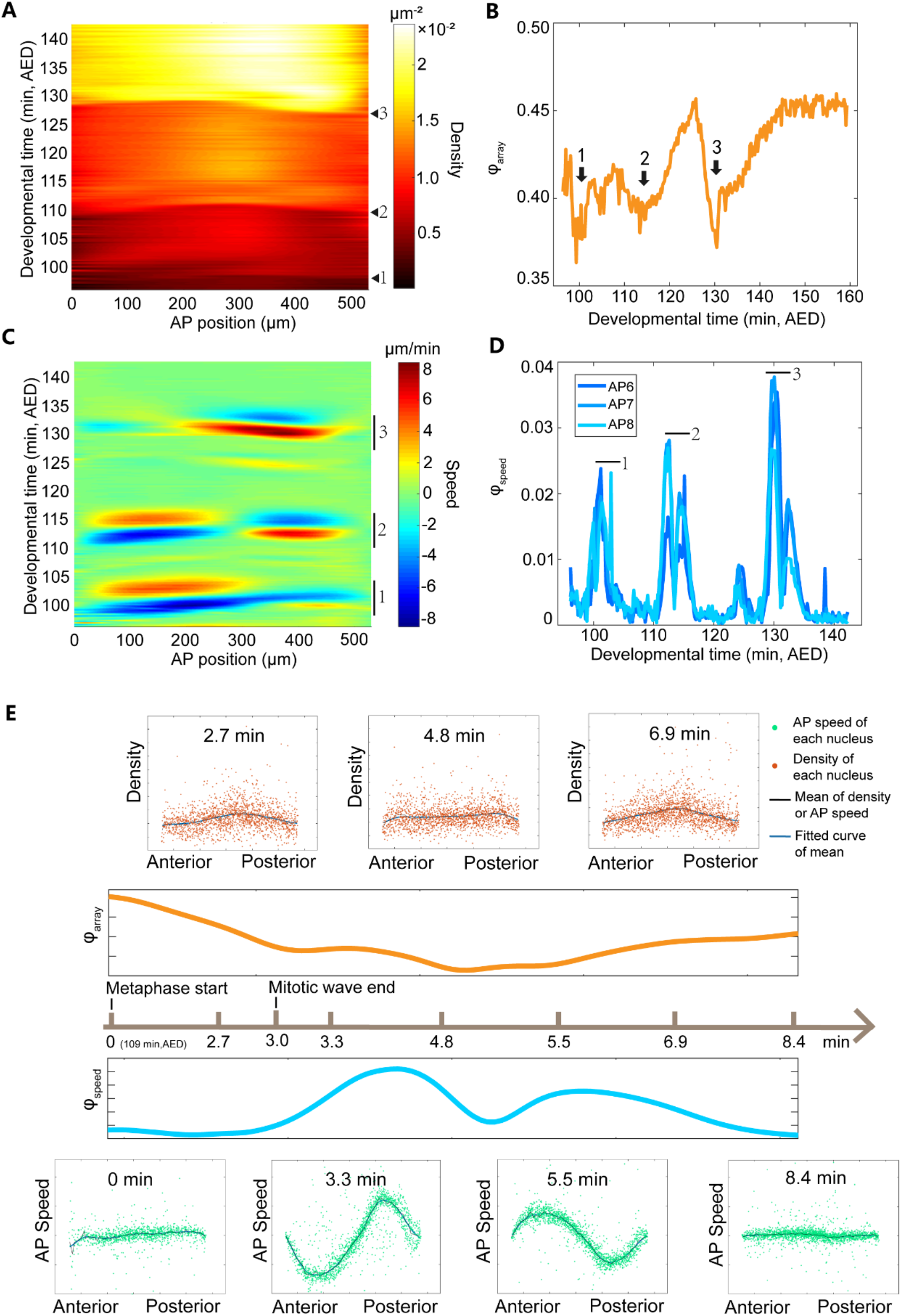
Characteristic features of the collective motion pattern and packing pattern of the nuclear array in *Drosophila* embryos from NC11 to NC14. (A) A representative heat map of the projected nuclear density along the AP axis of the embryo in each AP position from NC11 to NC14, i.e., rescaled developmental time from 90 min to 150 min after embryo deposition (AED) at 25 □ (see Supporting Information). Black triangles label the spindle split time points during mitotic phase 11, 12 and 13. (B) The dynamics of hexatic bond-orientational order parameter computed according to the equation 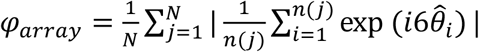. The black arrows 1, 2 and 3 label three time intervals around mitotic phase 11, 12 and 13 showing the minimum order parameter. (C) A representative heat map of the nuclear speed projected along the AP axis of the embryo in each AP position from NC11 to NC14. Positive and negative direction point to the posterior and anterior pole, respectively. (D) Dynamics of the order parameter of the collective motion from NC11 to NC14. The average order parameter of all the nuclei in each bin was computed according to the equation 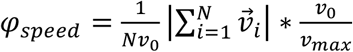. APi (*i*=6-8) corresponds to the *i^th^* bin with the width of 10% EL from the anterior pole (for the other bins, see Fig. S2). The markers 1, 2 and 3 label three regions that show high motion collectivity in (C) and (D). (E) The corresponding dynamics of the nuclear density, speed, smoothed *φ_array_* and smoothed *φ_speed_* of the nuclear array after the start of metaphase 12 shown in (A-D).

Intuitively, one might think that these nuclei are uniform-sized and arranged as hexagonal close packing, which is the closest packing on a plane. To our surprise, in the S phase from NC11 to early NC14 before cellularization, the hexatic bond-orientational order parameter (Kanesaki et al., 2011, Zippelius et al., 1980, Strandburg, 1988, Halperin and Nelson, 1978) increases from 0.35 to 0.45 (Fig. 2B and Fig. S5), significantly less than 1, the order parameter of a perfect hexagonal array. This result is consistent with a previous study showing that the proportion of the nuclei with six neighbors is approximately 56% (Kanesaki et al., 2011). The nuclear density also shows a non-uniform distribution along the AP axis. During the S phase of NC11 to NC14, the density ratio, which is defined as the ratio between the anterior (~0-5% EL) or posterior (~95-100% EL) density and the maximal density in the middle of the embryo during S phase, is about 0.65 (Fig. 4E). Hence the nuclear density is relatively 35% higher in the middle of the embryo than that in the pole region (Figs. 2A, S3, and 4E). And as the nuclear number doubles, the density ratio slightly increases (Fig. 4E), i.e., the nuclear distribution becomes more uniform. Previous results measured with fixed embryos also show higher nuclear density in the middle along the AP axis in NC14 (Hendriks et al., 2006, Keränen et al., 2006).

The density distribution and the regularity of the nuclear array are not always stable, they dynamically change as a function of the nuclear age *T* (defined as the time elapse after the metaphase, i.e., the start time point of spindle splitting). After *T*=0, the packing order of the nuclear array drops dramatically (Figs. 2B, 2E, and S5), then it is restored in approximately 8.4 min (Fig. 2E). Meanwhile, the density distribution unifies along the AP axis temporally when *T*=2.7-6.9 min (Fig. 2E).

As the nuclear packing pattern dynamically changes, a collective nuclear motion pattern emerges (Fig. 2E). Just after *T*=0, the nuclei move towards the embryo poles, and then partially move back to the original position based on the nuclear trajectory plot (Fig. S6). This phenomenon is consistent with the recently reported “yo-yo”-like movement (Lv et al., 2020). Interestingly, for each collective motion following with the mitosis wave, the nuclear speed projected to the AP axis shows one period of a 3-node standing wave (Movie S2). The period is approximately 9 min (Figs. 2C, 2E and S4). The 3 nodes locate at the two embryo poles and the embryo middle. The maximum nuclear displacement decreases from the positions of the crest to the node of the standing wave (Figs. S6 and S14B). Besides, the amplitude of this standing wave damps. For instance, during S phase 13 the maximum average nuclear speed reaches 8 μm/min in the first half period, then decreases by ~30% to 5 μm/min in the second half (Fig. 2C). If we calculate the wave peak ratio, which is defined as the ratio between the maximal wave crest of the second half period and the first half period of the AP speed standing wave (Fig. 4F), it is ~0.62 on average. As a result, the nuclear does not fully recover to the original AP position (Fig. 4F). Moreover, the motion collectivity (calculated with the order parameter 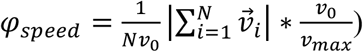 peaks during this collective motion process (Fig. 2D and Fig. S2).

The collective motion along the AP axis is significantly more pronounced than the one along the dorsal-ventral (DV) axis. Compared with the AP speed, the DV speed is very low (Fig. S1). Just after *T*=0, the nuclei slowly move to the dorsal side and then move back (Fig. S1). The maximum average DV speed is just 0.5 μm/min, much smaller than the maximum AP speed of 8 μm/min. And the DV speed is noisy. The absolute DV speed of each nucleus varies from 0 μm/min to 1.3 μm/min (Fig. S1A and movie S3). The standard deviation of the DV speed doubles during the collective motion process.

### Two spatio-temporal pulse-shape force fields learned with DNN show strong nuclear age dependence

To understand the origin of the observed nuclear packing and collective motion pattern, it is important to know the interaction rule between nuclei. The nuclei share a common cytoplasm from NC11 to early NC14, indicating they could share a common mechanism controlling the collective nuclear motion (Foe and Alberts, 1983, Farrell and O’Farrell, 2014). A reasonable assumption is that the spatio-temporal interaction rule between each pair of the nearest neighbor nuclei is identical. Here the spatio-temporal interaction rule is the internuclear force varying with space and time.

The motion of the nuclear array is controlled by the overdamped equation ignoring the inertia term and the internuclear friction term: 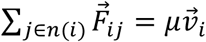. Where 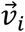 is the nuclear velocity, 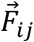 is the resultant internuclear force, *μ* is the viscosity of the cytoplasm (for details, see MATERIALS AND METHODS).

We assume that the absolute value of the internuclear force depends on the nuclear density and the nuclear age (the phase in a specific nuclear cycle). Because the nuclear speed is predominant along the AP axis (Figs. 2C and S1), the 3D single nuclear data is reduced to a 1D mean-field data (see MATERIALS AND METHODS). The single nuclear speed, density, and age are averaged in a bin of 5% EL along the AP axis (Fig. 3B). This data dimension reduction eliminates the single nuclear data noise and grasps the main features of the dataset, which is helpful for the later deep learning process.

**Fig. 3.**
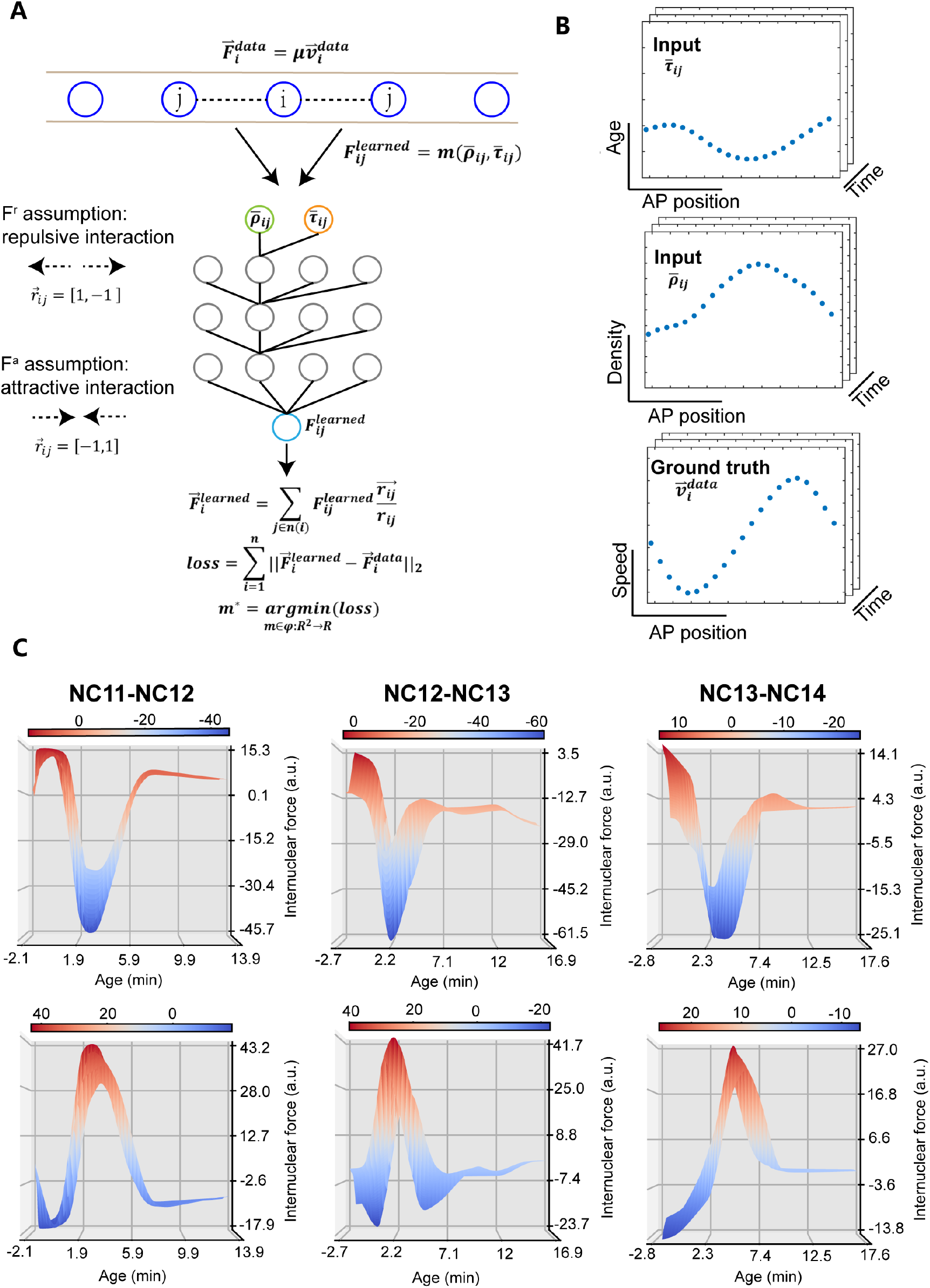
Force field functions learned from 1D data via MLFNN model. (A) The motion of nuclear array unit (*i^th^* unit) only interacts with the nearest neighbors (*j^th^* unit). The interaction force 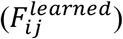 is the function of average nuclear density 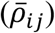 and average nuclear age 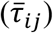 and 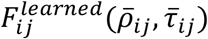 can be approximated by a three-layer MLFNN. The resultant force is calculated based on the assumed force unit vectors 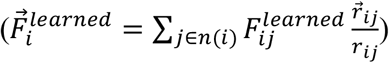. Based on F^r^ (or F^a^) assumption, 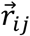 is different as shown in the figure. Then the learned resultant force 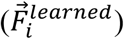 is compared to the ground truth 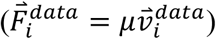 to define the loss function 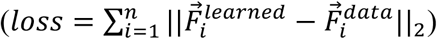 for training. (B) Snapshots of the training dataset. All the data are discretized along the AP axis. (C) The internuclear force has a conservative pulsating function relationship with the nuclear age, but has no clear correlation with the nuclear density. The top (bottom) panels are the DNN learning results based on F^r^ (F^a^) assumption. The labels “NC*-NC*” indicate the data used to be trained, e.g., “NC11-NC12” indicates the data from M phase 11 to S phase 12. Color bars show the magnitude of the force, and the positive or negative value of the force do not represent the force orientation.

It is convenient to learn the function form of *F_ij_* from the 1D nuclear array motion dataset (see MATERIALS AND METHODS) via a classical MLFNN (multilayer feedforward network) (Hornik et al., 1989, Svozil et al., 1997) model following the equation (Fig. 3A): 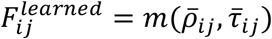. The input of the neural network is the average density 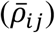 and average age 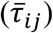 of the *i^th^* and the *j^th^* nuclear array unit. This can guarantee the interaction force between the two units is equal in the absolute value and opposite in direction. The output of the neural network is the average speed of the *i^th^* nuclear array unit 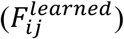 (Fig. 3A).

Since the interaction force *F_ij_* could be either attractive or repulsive, we take both possibilities into consideration. In brief, we call the repulsive force field as F^r^ and the attractive force field as F^a^. We define the direction towards the posterior pole to be the positive direction. The hyper-parameter force unit vectors for each pair of nuclear neighbor, representing the force orientation based on the F^r^ assumption and F^a^ assumption are 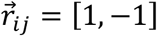 and 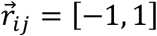, respectively. So the resultant force learned from the neural network is (Fig. 3A):

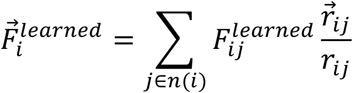

Because of the overdamped assumption, the ground truth of the resultant force is proportional to the motion speed (Fig. 3A):

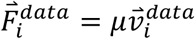

We search the best MLFNN model (*m\**) that fits the training data set (Fig. 3A):

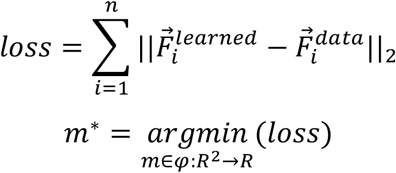

The training converges very quickly via the conventional back-propagation algorithm within 2000 training steps (Fig. S7).

The trained MLFNN models show that the absolute value of the internuclear force (*F_ij_*) strongly depends on the age of the nuclear array unit 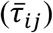, but not the density of the nuclear array unit 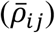. Based on the F^r^ assumption, *F_ij_* decreases along with 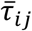 after *T*=0 and then increases to the original level (Fig. 3C). Based on the F^a^ assumption, the shape of the *F_ij_* function inversed (Fig. 3C). The force field forms from the MLFNN models in many different training trials are conservative except for some offset on the absolute value of the internuclear force (Figs. S8 and S9). That is because the relative internuclear force (resultant force) controls the motion pattern and no information about the initial value of internuclear force is added into the training process. Notably, color bars only show the magnitude of the force, and positive or negative values of the force do not represent the force direction (Figs. 3C, S8 and S9).

We confirm that the internuclear force form learned from MLFNN models is consistent with a deterministic mean-field physical model. Based on the overdamped momentum equation, we mathematically derive the force from the data (Supporting Information). We find that the internuclear force predominantly depends on the nuclear age and obeys a “negative-pulse-shape” curve with a width of around 10 minutes (Fig. S11C&D). Notably, here the positive or negative values of the force represent the force direction. The negative-pulse of the negative force actually represents the increase of the attractive force.

### A particle-based model using the F^a^ force assumption regenerates the packing pattern and collective motion pattern of the nuclear array

To test whether the force field learned from MLFNN models captures the observed characteristic collective behavior of the nuclear array, we run simulations with a coarse-grained particle-based model derived from the MLFNN models (see MATERIALS AND METHODS and Supporting Information).

Because 1D data was used to train the MLFNN model (Fig. 3B) and the motion is most pronounced along the AP axis (Figs. 2C and S1), we first run 1D simulations without the mitosis process. Consistent with the experimental results, the initial distribution of the nuclear array is sampled from the initial density distribution of S phase 13 in Fig. 2A. Using either the F^a^ or F^r^ force field (Fig. S12A), the heat maps of the AP speed and density match the experimental data (Figs. S12B&C, 2A&C and movie S4). However, the wave peak ratio equals one, i.e., the standing wave is symmetrical, so 1D simulation cannot recapitulate the phenomena of the damped standing wave or partial position recovery.

To closely mimic the biological condition, we then run 3D simulations on a prolate spheroid surface (see MATERIALS AND METHODS) and consider the mitosis process. Consistent with the embryo size, the ratio of the ellipsoid’s major axis and minor axis is 5:1.5 (Hendriks et al., 2006). In contrast to the 1D simulation, the number of the adjacent nuclei of one nucleus in 3D simulations is not constant. Considering the nuclear density is actually higher in the middle, the effective force between each nuclear pair should be less. Hence, the real internuclear force in 3D should be distance-dependent. And previous studies also show distance-dependent force fields are needed to stabilize the nuclear array on the ellipsoid surface (Dutta et al., 2019). And the distance-dependent force field usually has a “core region” and a “border region” (Dutta et al., 2019). Based on the F^a^ force assumption, the core region consists of the actin caps. To account for their specific sizes, a repulsive force is needed in the core region (Fig. S15A). And the actomyosin border (border region) acts like a spring (Zhang et al., 2018, di Pietro and Bellaïche, 2018), hence the attractive force increases along with distance (Fig. S15A). As for the F^r^ force, the motion of the nuclear array is mainly controlled by microtubules, so it is reasonable to assume the repulsive force decreasing along with distance in both the core region and the border region (Manhart et al., 2018, Kaiser et al., 2018, Mann and Wadsworth, 2019) (Fig. S15B). The age-dependent force field from MLFNN models is considered to describe the dynamics of the amplitude of the force (*A(T)*) in the border region, so only in this region in the simulation, the age-dependent force field is multiplied with the distance-dependent force field (Fig. S15). If we get rid of the core region in the F^a^ force assumption, the force field cannot maintain a regular nuclear array during S phase (Fig. S17).

The simulation results show that only the simulation based on the F^a^ force assumption recapitulates the characteristic features of the nuclear array. Just after *T*= 0, the nuclei migrate towards the embryo poles and then partially move back collectively (Fig. 4C&D and movie S5). The simulated AP trajectory recapitulates the experimental features (Fig. S14). The maximum nuclear displacement decreases from the positions of the crest to the node of the standing wave and the nuclei are not fully recovery to the original positions (Fig. S14A&B). Consistent with the previous experimental result (Lv et al., 2020), the simulation shows that faster mitotic waves associate with smaller maximum displacement (Fig. S14C). And the wave peak ratio in the simulation agrees with our experimental data (Fig. 4F). Moreover, the nuclear density stabilizes at a distribution that is higher in the middle and lower in the poles during S phase (Fig. 4A). And the density ratio of the simulation data is relatively lower than the median of all the experimental data but comparable with the experimental data of S phase 12 and S phase 13, during which the nuclear density is comparable to the simulation (Fig. 4E). The simulated order parameters of the nuclear array are also comparable with the experimental value during S phase (Figs. 4B and 2B). In contrast, based on the F^r^ force assumption, two extra collective motion processes show up before and after the experimentally observed motion process (Fig. S13C&D and movie S6). The nuclear array stabilizes at a nearly uniform density distribution during S phase (Fig. S13A). And the order parameters of the nuclear array are much higher than the experimental value during S phase (Fig. S13B).

**Fig. 4.**
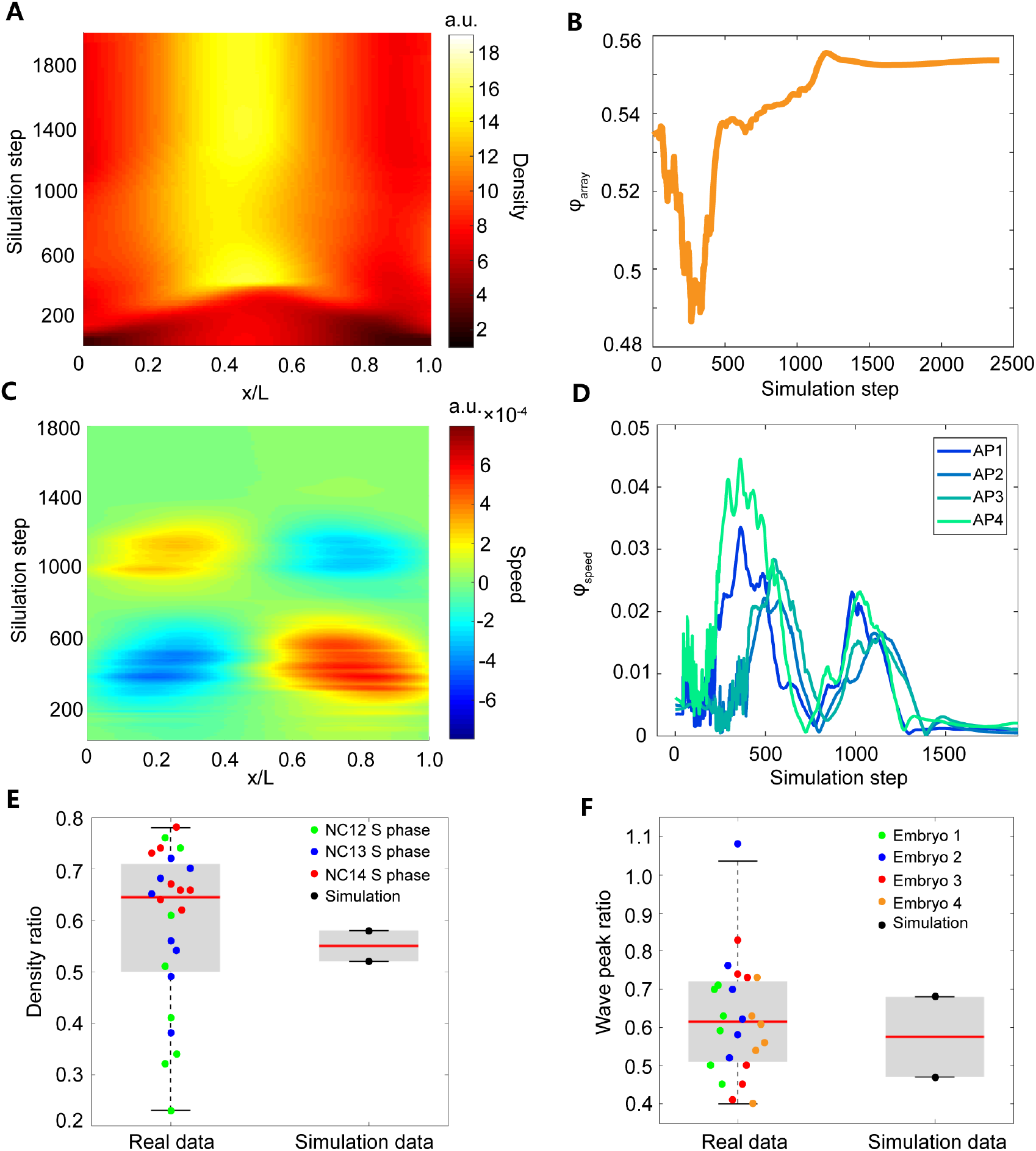
3D simulation results of the collective motion pattern of the wild type embryo based on the F^a^ force assumption (for the simulation movie, see movie S5). (A) Heat map of the nuclear density projected along the AP axis. (B) The dynamics of hexatic bond-orientational order parameter, which was computed by the equation 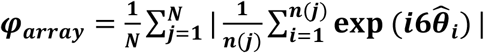. (C) Heat map of the nuclear speed projected along the AP axis. (D) Dynamics of the order parameter of the collective motion, which was calculated according to the equation 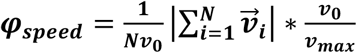. APi (*i*=1-4) corresponds to the *i^th^* bin with the width of 25% EL from the anterior pole (E) Density ratio is defined as the ratio between the anterior (~0-5% EL) or posterior (~95-100% EL) density and the maximal density in the middle of the embryo during S phase. The medians (red line) of measured and simulated density ratio are 0.65 and 0.55, respectively. (F) Wave peak ratio is defined as the ratio between the maximal wave crest of the second half period and the first half period of the AP speed standing wave. The medians (red line) of measured and simulated wave peak ratio are 0.62 and 0.58, respectively. The force field used in this simulation is shown in S15A.

The age difference along the AP axis originated from the mitotic wave is the key to generate the asymmetric force driving the directional collective motion of the nuclear array along the AP axis. Consider a nucleus in the first half period of the standing wave, the amplitude of the attractive (repulsive) force enforced from the embryo pole side is greater (smaller) than that from the embryo middle side due to the age difference, the resultant net force is towards the pole direction, hence the nuclei collective moves towards the pole. In the second half period, the amplitude of the attractive (repulsive) force enforced from pole side is smaller (greater) than that from the middle side, this results in a collective movement away from the pole (Fig. 6A).

However, only the distance-dependent attractive force is sufficient in maintaining the non-uniform density distribution. The nuclear density is higher in the middle than in the pole region. Hence the internuclear distance *r* is smaller in the middle. For the attractive force, it increases as *r* increases. For a nucleus under the nuclear density gradient, the individual pair-wise internuclear force is stronger from the embryo pole side, but the number of the interaction nuclei is more from the embryo middle side, the two factors could compensate each other to achieve a balance. In contrast, the repulsive force, which decreases as *r* increases, is always weaker from the pole side. As the nuclei close to the middle region have more neighboring nuclei with stronger repulsive force, they can push those close to the pole region (which have less neighboring nuclei with weaker repulsive force), hence the nuclei in the middle tend to move to the pole region (Fig. 6B).

Moreover, only the attractive force is able to replicate the observed collective motion. In the simulation with the F^a^ force field, the internuclear force act as an attractive force, so the newly divided nuclear regions tend to contract to pull metaphase nuclei to the poles of the embryo after *T*=0. And because the attractive force can stabilize the nuclear array with a higher density distribution in the middle of the embryo, no extra motion emerges (Fig. 4A&C). By contrast, in the simulation with the F^r^ force field, the internuclear force acts as a repulsive force, so the newly divided nuclear regions (near poles) with smaller *r* tend to expand after metaphase. This tissue expansion pushes metaphase nuclei to the middle of the embryo, which forms the extra motion process over-compressing nuclei in the middle during metaphase. And after a period of the standing wave, the higher density distribution in the middle of the embryo cannot be stabilized. The second extra motion process emerges to reduce the high density in the middle region (Fig. S13A&C).

The distance-dependence of the attractive force implemented in 3D simulations is the key to recapitulate the experimental wave peak ratio (i.e., the dampening standing wave) and partial position recovery in the simulation. In 1D simulations, the relative higher nuclear density distribution can only be maintained if the force is distance-independent. For the internuclear force only depending on the nuclear age, and the nuclear age uniformly increases among all the nuclei, so the resultant force is symmetric in two half periods of the standing wave (Fig. S12C). By contrast, in 3D simulations based on the F^a^ force assumption (Fig. S15A), the age-dependent attractive force also increases at larger internuclear distance *r*. Since the embryo shows an initial higher nuclear density and younger nuclear age in the middle region, the resultant net effective force is asymmetric in two half periods. In the first half period, the attractive force shows larger amplitude *A(T)* and *r* from the pole side, and smaller *A(T)* and *r* from the middle side. But in the second half period, the pole side changes to smaller *A(T)* and larger *r*, and the middle side changes to larger *A(T)* and smaller *r*. Hence the resultant force from the first half period is greater than the one in the second half (Fig. 6C).

### The prediction based on the F^a^ force field function is confirmed with the observed nuclear motion patterns

To further test the F^a^ force field learned from MLFNN, we apply it in three extra conditions, in which the experimental observation has not been explicitly utilized in the target function in the model training.

In the training dataset shown as the collective motion after the M phase 12 in Fig. 2C, the nuclear position with the maximum speed, i.e., the anti-node position of the standing wave of the AP speed, is at 25% EL from the embryo pole. This appears to be a coincidence as it only shows up when the two mitosis waves from the two poles nearly synchronize. As we vary the time difference of the two mitosis waves, the anti-node position in the middle also shifts linearly away from the pole where the leading wave initiates (Fig. 5A&C). Indeed, in the experiment, we also observe a linear change of the middle anti-node position from 30% EL to 70% EL as the start time of the mitosis wave from the two poles differs from 0 min to 3 min (Fig. 5A). And the density ratio has no significant change compared to the synchronized division case (Figs. 5B and 4E).

**Fig. 5.**
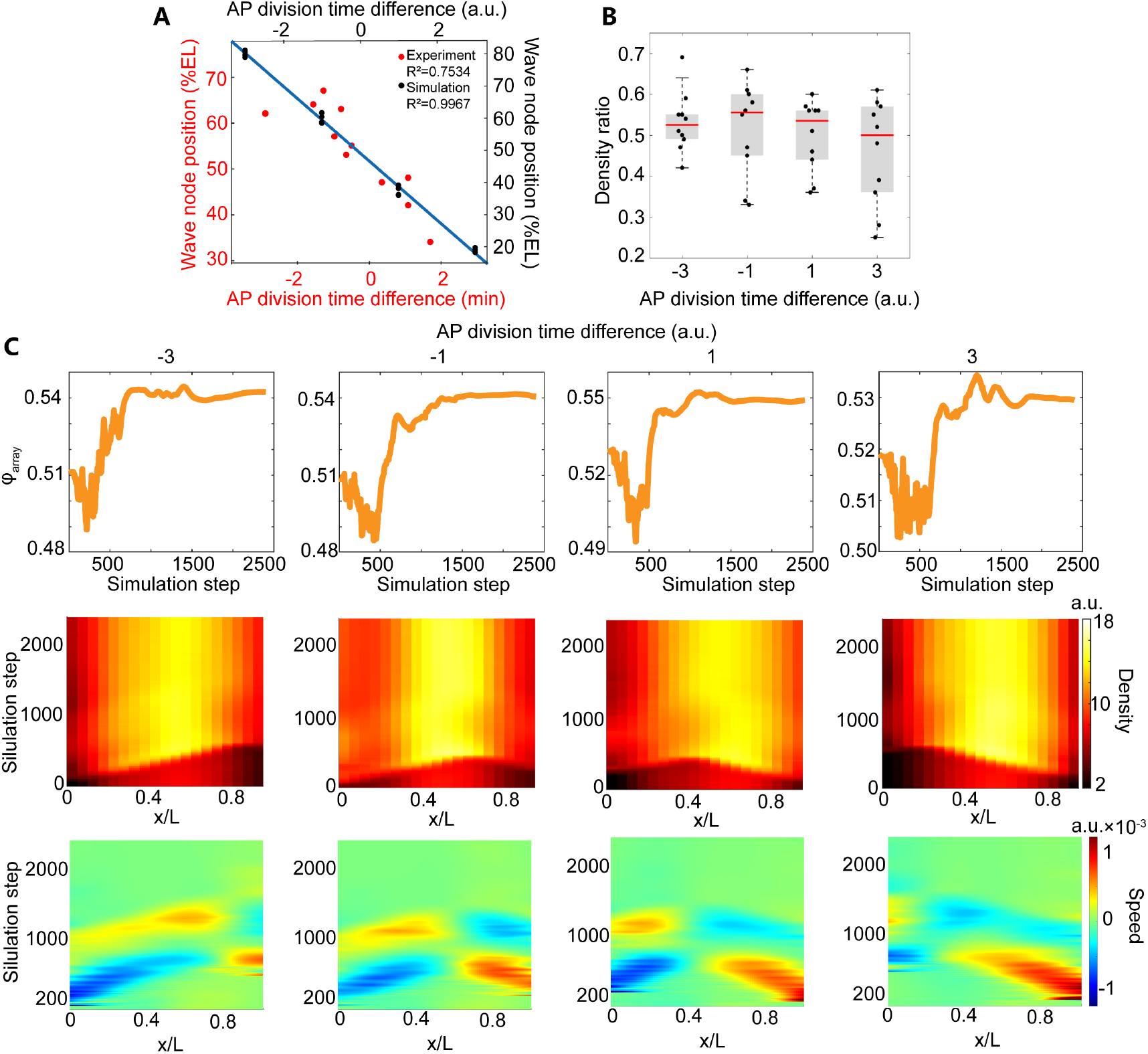
3D simulation results of the embryos with varied time difference between the start of metaphase at two poles (F^a^ force assumption). (A) The linear relation between the position of the second node of the standing wave of AP nuclear speed and the division time difference between the anterior and posterior poles of experimental (red) and simulation (black) data. (B) The density ratio of the simulation data with different AP division time difference. The medians of the four group data in the panel are 0.5, 0.535, 0.555 and 0.525, respectively. (C) The representative characteristic features of the collective motion pattern and packing pattern of the nuclear array in the simulations with different AP division time difference. The force field used in this simulation is shown in S15A Fig.

**Fig. 6.**
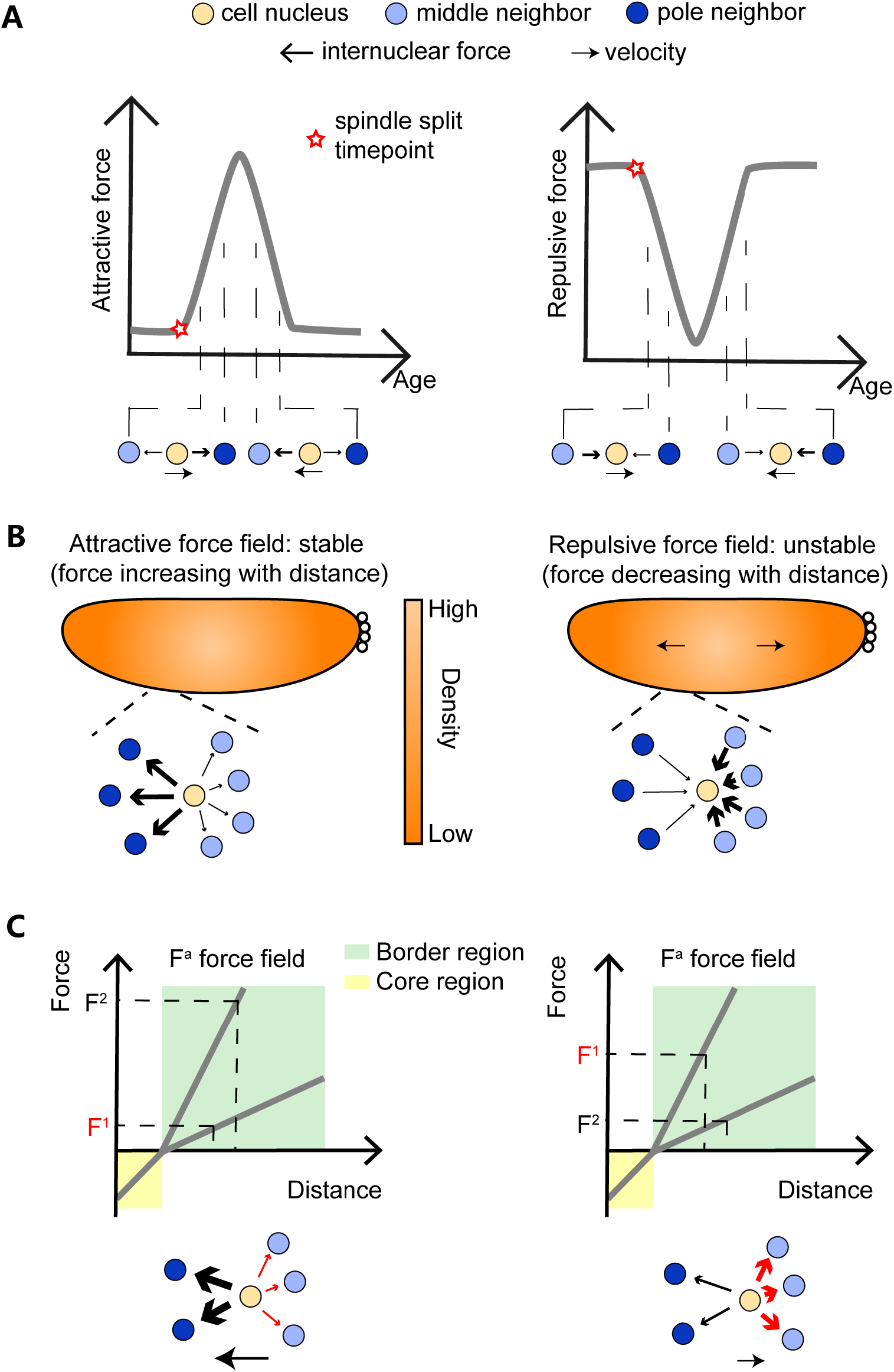
The intrinsic properties of the force fields generate the characteristic features of the nuclear array. (A) Collective motion is driven by the age-dependent force field. (B) Density asymmetric distribution is maintained by the attractive distance-dependent force field. (C) The dampening standing wave of the nuclear speed is derived from the attractive force field with the dependence of the nuclear age and distance.

In the extreme case, the mitotic wave could reach the other pole before the initiation of the opposite mitotic wave. We simulate this condition and find that the standing wave of the AP speed changes to be two-node (Fig. S16A). But the final density distribution after mitosis and the dynamics of the pack order are still similar to the one with two mitotic waves (Fig. S16A). This condition actually exists in a small number of wild type embryos. And our simulation results are consistent with the experimental results observed in these embryos (Tomer et al., 2012).

The standing wave of the AP speed could further be adjusted to be 5-node if fastening the embryo in the middle. Our simulation is consistent with the experiment (Deneke et al., 2016). Again this does not change the density distribution during S phase and the level of nuclear array regularity compared with wild type embryos (Fig. S16B).

## DISCUSSION

Light sheet microcopy helps to reveal the collective packing and motion pattern of the nuclear array in *Drosophila* embryos. As an excellent model system of collective motion, the nuclear array in *Drosophila* embryos has already been reported in several studies. Nearly all the previous experimental data were obtained with confocal microscopy. Due to the limited imaging speed, the embryos were often softly pressed to generate a flat surface filled with nuclei in a shallow depth. It has been reported that the press on C. elegans embryos could induce profound change in cell movement during embryogenesis (Jelier et al., 2016, Giurumescu et al., 2012). And it has also been proposed that the mitosis wave in *Drosophila* embryos is a mechanical wave (Idema et al., 2013), hence, the movement of the nuclear array could also be affected by the mechanical stress in pressing embryos. Here we use a light sheet microscope for 3D imaging. Instead of imaging only several micrometers in depth below the flattened embryo surface with confocal microscopy, we could image nearly the whole embryo with the comparable overall time resolution. Moreover, phototoxicity is much lower for longer live imaging time. Most importantly, benefitting from the sample preparation method (embedding the embryo in agarose), the embryo has no deformation and the natural membrane curvature is maintained. This leads us to find a stereotypical density distribution along the AP axis: the nuclear density is relatively 35% higher in the middle of the embryos than that in the pole region during the S phase of NC11-NC14 (Fig. 4E). And the similar result in NC14 has only been reported in a previous measurement on the whole fixed embryos with two-photon microscopy (Hendriks et al., 2006, Keränen et al., 2006). These results suggest that the nuclear density may be influenced by the membrane curvature and it is important to keep the embryo shape intact during imaging. We also discover that the collective motion of the nuclear array shows a 3-node standing wave in AP speed for the first time.

Reversal engineering with DNNs successfully extracts the net force field governing the collective behavior of the nuclear array in *Drosophila* embryos. Although several particle-based models for the nuclear array have been reported, none of them can account for the nuclear collective motion pattern discovered in our study. For example, the model in ref. (Dutta et al., 2019) cannot generate any collective motion at all. The model in ref. (Koke et al., 2014) generates a collective motion with the opposite direction. The model in ref. (Lv et al., 2020) shows similar results as our repulsive force simulation results, an extra motion appearing after the normal motion process. Instead of giving an ad hoc formula of the force field, we reversely extract the net force field using MLFNN model. We find that the resultant F^a^ force field learned from MLFNN model show strong nuclear age dependence. To stabilize the 3D simulation, we further multiply F^a^ with a distance-dependent force to account for the “core-border” structure of the nuclei. Instead of using a simple sphere surface, we run a 3D simulation on a prolate spheroid surface, which mimics the embryo shape better. The simulation results are consistent with all the observation in the experiment. Hence, this force field is the most likely force field implemented in the embryo. And the DNN-based method demonstrated in our study could be a powerful tool in extracting the force field underlies the other complex collective behavior.

However, what biological mechanism generates this force field remains to be discussed. Based on our results and previous studies (Zhang et al., 2018, Royou et al., 2002), we can speculate a mechanism as following. Right after metaphase, the mitotic furrow recovers and the older nuclei group (close to the pole regions) forms actomyosin borders between each nucleus. The attractive force provided by the newly formed actomyosin borders pulls the tissue away from the mitotic wavefront. Note that, metaphase nuclear region is being pulled by the anaphase region (myosin enrichment region), which is consistent with the previous study (Lv et al., 2020). And when the M phase is finished, the younger nuclei group has a larger internuclear distance (membrane deformation) than the older nuclei group because of the previous pulling process. As a result, the tension is larger between the younger nuclei than the older nuclei. Hence, the nucleus array moves back because of the tension difference (Fig. 7A). Several lines of experimental results seem to support this speculated mechanism. For example, the cell deformation is larger in the middle of the embryo than the pole region (Fig. 7B). And the nuclear collective motion pattern nearly disappears after injecting myosin II inhibitor into the embryo (Lv et al., 2020), indicating that myosin II may be a core upstream factor to generate the collective motion. Further experiments are still needed to test this speculation directly.

**Fig. 7.**
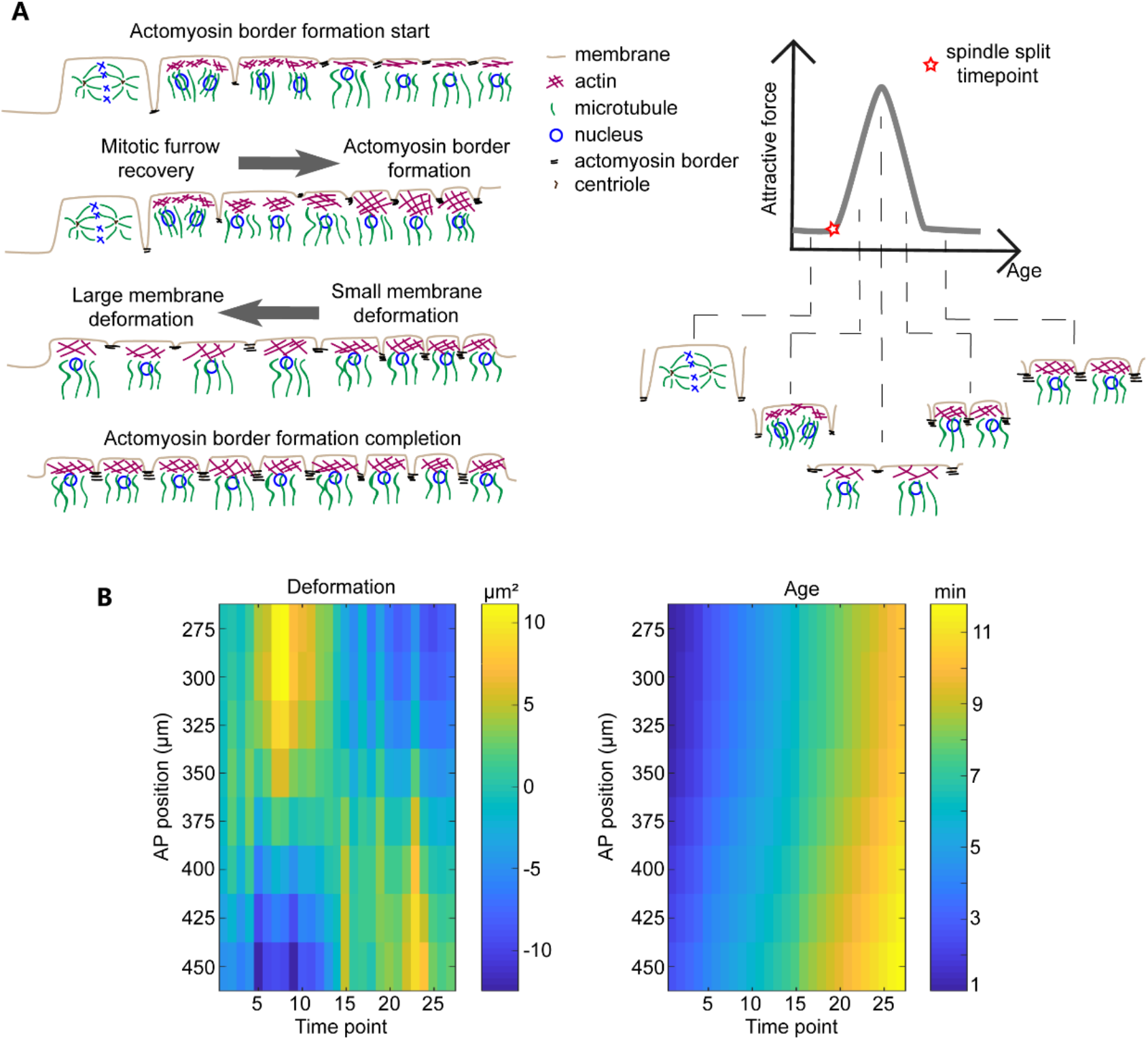
Molecular basis of the F^a^ force field from the MFLNN model. (A) The relationship between biological molecular dynamics and the F^a^ force. Each nucleus dynamically changes between five typical states during each nuclear cycle: “mitotic furrow state”, “mitotic furrow recovery state”, “flat membrane with lager membrane deformation and less myosin II state”, “actomyosin border formation with small membrane deformation state”, and “actomyosin border formation completion state”. The relationship between the states and the F^a^ force field is shown in the right panel. The corresponding nuclear motion process is shown in the left panel. (B) Heat maps of the calculated average cell area (the inverse of nucleus array density) variation relative to the original cell area (left) and the nuclear age (right) during S phase 13.

The precise and reproducible packing of the nuclear array should be crucial in controlling the cell size, establishing developmental patterns, and coordinating morphogenesis in fly embryos. Most of the previous studies focus on the packing patterns of the nuclear array. During S phase, based on the radial distribution function data, the nuclear array has no long-range positional order, nor is amorphous (Dutta et al., 2019). And based on the KTHNY theory (Zippelius et al., 1980, Strandburg, 1988, Halperin and Nelson, 1978), during S phase the nuclear array is the liquid state because of the topological defects (5 nuclear neighbors or 6 nuclear neighbors) (Kanesaki et al., 2011) and exponential decay of the hexatic correlation function (Kaiser et al., 2018). The nuclear density distribution remains high in the middle of the embryo from NC11 to NC14. On the one hand, such a distribution is robust and it could be maintained as long as collective nuclear motion is generated based on the F^a^ assumption. On the other hand, such a density distribution may also help to maintain the collective motion. Assuming the mitosis wave is triggered at the locations with a low nuclear density, the mitosis wave will always start from the two poles and move to the middle of the embryo. The nuclear age difference along the AP axis leads to the asymmetric force driving the collective nuclear motion along the AP axis. This density distribution could be generated by the self-organized nuclear spreading from the middle to the whole embryo during the pre-blastoderm (Deneke et al., 2019). Hence the nuclear packing pattern and the collective motion pattern consist a self-sustainable loop. The biological significance of these collective behaviors of the nuclear array remains to be explored in future studies.

## MATERIALS AND METHODS

### 4D live imaging of early *Drosophila* embryos

#### *Drosophila* embryo sample preparation

The fly stock was maintained at 25°C on cornmeal medium. His-GFP fly line (from Thomas Gregor Lab at Princeton University) embryos were collected in 1.5 hours on the grape juice plate, then dechorionated in 4% NaClO bleach buffer for 2 min. After being washed in ddH_2_O for several times, the embryos were mounted in 1% low melting agarose in the capillary. For the convenience of imaging, the embryos were carefully mounted along the AP axis and the light sheet can transmit from the dorsal or ventral side of the embryo.

#### Light sheet microscope imaging

Imaging experiments were performed with a Zeiss Light Sheet Z.1 Microscope The illumination objective is LSFM 10×/0.2, and the detection objective is W Plan-Apochromat 20x/1.0 Corr DIC M27 75 mm. Under DIC imaging, the pole cell formation process can be clearly identified, which is a marker for the start of NC10. Starting from NC10, the imaging lasts for about 2 hours to cover four mitotic phases from NC10 to NC14. H2A-GFP is excited by a 488 nm laser with 1~2% laser intensity and the exposure time of 30 ms to avoid high phototoxicity which may cause nuclei falling to the yolk (Kanesaki et al., 2011). The emission light from 505 nm to 545 nm was collected. To study the motion pattern, a relatively high imaging speed is needed. The *z* stack images (1920×1920 pixels, pixel size 0.286 μm) were acquired in 1 μm steps at 20 s time intervals. To achieve high temporal resolution, the embryos were only imaged from one angle, so that three quarters of the embryos were captured. “Dual Side when Experiment” mode and “Pivot Scan Settings” mode were selected while imaging to achieve dual side illumination and reduce shadows which might be cast by optically dense structures within the sample.

The protocols from the ref. (Yue et al., 2020) was applied in the nuclear segmentation process.

### Data analysis

#### Projected nuclear speed, trajectory and density along the AP axis

After nucleus segmentation process, the 4D nuclear position information was obtained. At each time point, the nuclear coordinates are a collection of data points on the embryo surface, i.e. a point cloud. The middle points of scattered cell nuclear point cloud in the anterior end and posterior end were manually picked as the anterior pole and posterior pole.

The nuclear position *x*(*t*) was projected to the AP axis to obtain the AP projected trajectory. The nuclear speed was calculated according to the formula *v*(*t*) = (*x*(*t*) – *x*(*t* – Δ*t*))/Δ*t*. Here Δ*t* is the imaging time interval, i.e., 20 s. For plotting the heatmap of the nuclear speed, in each time point the nuclear speed was averaged in each bin with the width of 10% EL along the AP axis.

The embryo surface was reconstructed from the scattered cell nuclear point cloud with the function MyCrustOpen.m (from MathWorks File Exchange website: https://ww2.mathworks.cn/matlabcentral/fileexchange/63731-surface-reconstruction-from-scattered-points-cloud-open-surfaces). After reconstruction, the point cloud was triangulated. Using Voronoi diagram, each triangular facet can be divided into three quadrangles. Each quadrangle area contains a triangle vertex (a cell nucleus). The nuclear density was obtained by taking the inverse of all the quadrangle area. For plotting the heat map of the nuclear density, in each time point the nuclear density was averaged in each bin with the width of 10% EL along the AP axis.

#### Order Parameters

The most efficient and the closest packing on the plane is hexagonal close-packing. To evaluate the regularity of the nuclear array in the *Drosophila* embryo, the hexatic bond-orientational order parameter is calculated with the formula 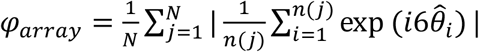 (Kanesaki et al., 2011). *N* is the total number of the nuclei, *n* is the neighbor number of each nucleus and *θ* is the angle between vertical vector and the vector between each nucleus (i^th^ nucleus) and its neighbor nucleus (j^th^ nucleus) (Kanesaki et al., 2011). The value of *ψ_array_* varies from 0 to 1. The larger the value, the closer the nuclear array to the hexagonal closed packing array.

To evaluate the degree of nuclear array motion collectivity during the developmental process in the *Drosophila* embryo, the order parameter in the field of collective motion defined by the equation 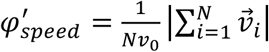, is used(Vicsek and Zafeiris, 2012). *N* is the total nuclear number, *v*_0_ is the average absolute nuclear velocity. The value of *φ_speed_* also varies from 0 to 1. If the motion is disordered, the velocities of each nucleus point to random directions and average out to give a small magnitude vector, whereas for ordered motion the velocities add up to a vector of absolute velocity close to *Nv*_0_. To account for the potential effect of the velocity difference, the equation is amended to 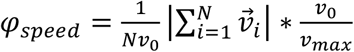. Here *v_max_* is the maximal velocity of all the nuclei in all time points.

### Learn the internuclear force field function from data via a multi-layer feedforward neural network

#### One dimensional nuclear array motion dataset

In the 1D condition, a single nucleus is no longer a motion unit. Instead, all of the single nuclear data including nuclear density, nuclear age and nuclear speed are averaged along the AP axis in each time point and fitted with a smooth spline. Then the 1D discrete data is obtained from the spline in 25 μm interval along the AP axis (Fig. 3B). Here, each position along the AP aixs is a nuclear array unit. And the corresponding density, age and speed represent the average value of the nuclei around this AP position. Then in the 1D condition, a nuclear array unit is a new motion unit. Note that, the two data points in the embryo poles in each time point are considered to be fixed and not included in the dataset.

All the input data including nuclear density data and nuclear age data is normalized by the equation 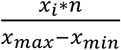. Here, *n* equals 100. After normalization, the two datasets have similar data range, which ensures that the two datasets have similar contribution for training the DNN.

Note that, the start points of the nuclear cycle are calculated from the density gradient data (Fig. S10).

#### DNN architecture and loss function

The multi-layer feedforward neural network used here has 3 hidden layers, each with 100 nodes. The input layer has 2 nodes and the output layer has only 1 node. Softplus function (*f*(*x*) = *log*(*e^x^* + 1)) is used as the activation function except for the output layer because the output value is not always being positive value.

The loss function is defined as the Euclidean distance between the experimental data 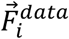 and the training results 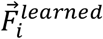 of all cell nuclear array units.

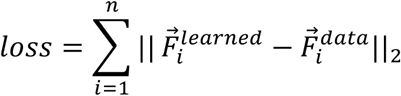

#### Training

The DNN is trained by Adam optimizer in tensorflow. The learning rate is set by the function tf.train.exponential_decay. The initial learning rate is 0.05 and exponentially decay in each 500 steps with the decay rate 0.96:

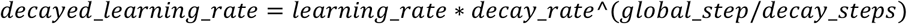

1D nuclear array motion dataset is used to train the DNN firstly. Note that, the data in the training dataset is sparse and discrete but the trained DNN can be used to explore a much larger parametric space. To get a continuous internuclear force field function from the trained DNN, a new input whose data range is the same with the training dataset but has a really small data interval (only 0.1) is feed to the DNN. The new input and the corresponding DNN output is plotted in a three dimensional space as the internuclear force field function.

The training results are not sensitive to the width of hidden layer and depth of the neural network. The DNNs have two or more hidden layers each with ten or more nodes can give out similar force fields using the same training data set. The details of the DNN architecture used in the main text are shown in the “DNN architecture and loss function” part.

### The particle-based model to simulate nucleus array collective motion pattern on the prolate spheroid surface

#### The basic particle-based model

Strictly speaking, the motion of the nucleus array should be described by the following equation, which contains an inertia term, a pressure term, a cytoplasm friction term and an internuclear friction term (for simplicity, here is the one dimensional equation):

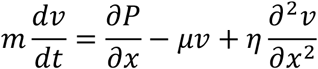

, where *m* is the mass of the nucleus, *v* is the velocity of the nucleus, *P* is the internucleus pressure, *μ* is the viscosity of the cytoplasm, *η* is the internuclear friction. The internucleus pressure is mainly controlled by actin, myosin II and membrane structure (Zhang et al., 2018, di Pietro and Bellaïche, 2018, Schmidt and Grosshans, 2018, Foe et al., 2000). The internuclear friction is mainly controlled by microtubules (Koke et al., 2014).

Inside the embryo, Reynolds number is really small (*Re* ≈ 10^−5^) so that nuclear motion is dominated by viscous forces and inertia term can be omitted (Deneke et al., 2019). Because collective motion order of the nuclear array is high during early S phase and nuclear array is relatively stable during late S phase (Fig. 2C), 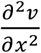 is relatively small, so that we also omitted the internuclear friction term. Then the function is simplified to the overdamped equation:

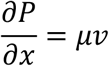

The discretization version of the overdamped equation in three dimension is:

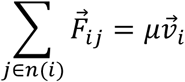

, where 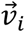 is the velocity of the nucleus, 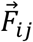 is the resultant internucleus force, and *μ* is the viscosity of the cytoplasm.

The overdamped equation of motion is also used in the following biological systems: muscle cells nucleus positioning (Manhart et al., 2018), *Drosophila* embryo nucleus array packing and motion (Dutta et al., 2019, Lv et al., 2020, Kaiser et al., 2018), and *Caenorhabditis elegans* embryo cell positioning (Fickentscher et al., 2013, Yamamoto and Kimura, 2017).

While simulation, the initial nuclear number is 400. After mitosis, the nuclear number doubles. The nuclear number of the simulation is comparable with the nuclear number in the S phase from NC10 to NC11. In the prolate-spheroidal coordinate system, the initial nuclear coordinates (*v*, *θ*, *φ*) are set as the following: *v* is a constant, *θ* is sampled from Gaussian distribution, and *φ* is sampled from the Uniform distribution. Because after nuclei migrating from yolk to the cortex region during the S phase of NC10, the initial nuclear density is higher in the middle than the pole regions in the embryo (Deneke et al., 2019), it is reasonable to use Gaussian distribution of *θ* to imitate this process. Then the random nuclear coordinates are rearranged under the distance dependent force fields to form a new nuclear array as the initial state.

In wild type embryos, mitotic waves usually start from the anterior pole and posterior pole (Deneke et al., 2016, Vergassola et al., 2018). That means the nuclear age has a slight phase difference. The initial nuclear age distribution is shown in Fig. S15C. The nuclear division orientation is random (Lv et al., 2020) in simulations. After metaphase, the internuclear distance between daughter cells is within the core region of the distance-dependent force fields, so that daughter cells are repulsive to each other and no additional “nuclear division repulsive force” is added to the force field.

For the equations of motion in polar coordinates on the prolate spheroid surface, see supporting information.

## Acknowledgement

We would like to thank Xiang Liu and Jingxiang Shen for helpful discussions; Jingyi Wu for the help of nuclear segmentation and National Center for Protein Sciences at Peking University for Zeiss Light Sheet Z.1 Microscope.

## Author Contributions

X.X.W. and F.L. designed the study. X.X.W. performed the experiments and analyzed the data. X.X.W. and K.K.K. developed the models. W.L.X. performed the image processing. X.X.W. and F.L. wrote the paper with the input from W.L.X. and K.K.K.

